# Natural variation of gibberellin levels in maize tissues across North America

**DOI:** 10.64898/2025.12.18.695287

**Authors:** Xia Guan, Nitin Supekar, Changjian Jiang, Junhong Guo, Edwards Allen, Charles Dietrich, Cody Postin, Yanfei Wang

## Abstract

Gibberellins (GA) are endogenous phytohormones that regulate growth and development throughout the plant lifecycle. While GA has been extensively studied under controlled conditions for their fundamental role in plant adaptation and agronomic trait development, field-based data on their natural variation across diverse agricultural genotypes and environments remains limited. To address this knowledge gap, we analyzed bioactive gibberellin A1 (GA_1_) and gibberellin A4 (GA_4_) across multiple maize hybrids grown in diverse field environments using high-throughput liquid chromatography-tandem mass spectrometry (LC-MS/MS). This study provides comprehensive quantitation of GA_1_ and GA_4_ levels in maize tissues across diverse commercial hybrids grown under standard agricultural practices, revealing hormonal variation due to genetic background and environmental conditions. Our findings demonstrate that GA_1_ and GA_4_ concentrations from conventional maize samples varied across tissue types and field sites. Analysis of variance components indicated that the site variation was statistically significant at the 5% significance level in five out of ten GA and tissue combinations, while the genetic component was significant only for GA_4_ in grain samples. Notably, significant interaction between hybrid and site was observed for both GA_1_ and GA_4_ in stalk tissue. Positive correlations between GA_1_ and GA_4_ were observed consistently across tissues except root. These results underscore the common metabolic pathways involved in regulating GA levels in maize and provide the first comprehensive dataset of GA_1_ and GA_4_ variation under real-world agricultural conditions. Our findings offer insights into the interplay between genetics, environment, and hormonal regulation in maize and suggest potential targets for breeding programs aimed at improving agronomic traits such as yield.

## 1. Introduction

Gibberellins (GA) are endogenous phytohormones that play a crucial role in regulating plant growth and development throughout the plant lifecycle. They were first discovered as secondary metabolites in the fungus *Gibberella fujikuroi*.[1, 2] GA comprises a class of diterpenoid carboxylic acids that possess the tetracyclic ent-gibberellane carbon skeletal structure. Among various GA, only a few, e.g., GA_1_, GA_3_, GA_4_ and GA_7_ exhibit significant biological activity[3]. GA_1_ and GA_4_ are the major active GA that promote growth and development in plant species. They regulate and control diverse developmental processes such as seed germination, stem elongation, leaf expansion, and flower and fruit development[2]. Additionally, GA regulates plant adaptation to both biotic and abiotic stresses[4, 5].

The biosynthesis and signaling pathways of GA are intricate systems that exhibit complex spatiotemporal patterns across cells, tissues, and developmental stages [6]. These patterns result from multiple enzymatic steps and feedback mechanisms that respond dynamically to both endogenous and environmental signals [7, 8]. Environmental factors such as drought stress can significantly reduce GA biosynthesis across plant species[9], leading to stunted growth and altered development, while optimal conditions enhance GA production and promote vigorous growth[10]. The movement of GA between cells and across plant organs, coupled with their variable distribution patterns, creates challenges for understanding hormone-environment interactions in agricultural settings[11].

Due to the vital biological activity of GA in plant growth and development, several applications have been developed with commercial and biotechnological utilization: exogenous applications of GA to promote growth, and exogenous applications of its concentration or the genetic modification of biosynthesis to optimize plant phenotypes and environmental responses[8]. Research in recent years has shown that GA have direct and indirect effects on the regulation of many plant traits. Many plant varieties with improved agronomic traits (e.g., dwarf phenotypes or increased biomass) are shown to be related to GA activity and signaling[12], thus manipulation of GA levels is commonly used in agricultural practices to optimize plant growth and yield.

Maize (*Zea mays* L.; Corn) is a major agricultural commodity, contributing significantly to the economy of many countries and serves as a staple food for many populations worldwide[13]. Short maize hybrids have been created through conventional breeding by researchers with the objectives to improve performance under high density and improve lodging tolerance[14]. However, breeding short traits into existing maize hybrids is time consuming, challenging, and can affect environmental responses. Genetically modified maize can be engineered through the enhancement or suppression of hormone pathways including gibberellin biosynthesis. Scientists have developed a genetically modified short stature maize ideotype by suppressing maize genes that regulate the levels of GA, reducing plant stature without undermining maize yields[15].

Accurate quantification of GA in plant tissues has been instrumental in understanding GA-mediated mechanisms and effectively improving their practical applications[16]. While previous studies have quantified bioactive GA in maize tissues[15, 17–21], these analyses were typically limited to specific varieties or locations, lacking the scope needed to characterize natural variation across diverse agricultural conditions. However, establishing baseline patterns of GA variation in agricultural settings is crucial for several reasons. First, understanding the natural variability of GA concentrations in conventional maize tissues - a primary determinant of plant stature – and distinguishing between environmentally-induced and genetically-controlled variation provides essential context for crop improvement efforts. This baseline knowledge is particularly valuable for developing and evaluating new short-stature varieties, where GA-related genetic modifications must be assessed against the background of natural GA fluctuations that occur across diverse field environments.

Second, while GA biology has been extensively studied under controlled conditions[15], quantifying the magnitude of GA variation in commercial hybrids under real-world agricultural conditions helps bridge the gap between laboratory findings and field performance. This knowledge is critical for predicting the stability and effectiveness of GA-modified traits across different growing environments, where environmental stresses can significantly impact GA-mediated growth processes and ultimately affect yield potential. Understanding the relative contributions of genetic background and environmental factors to GA concentrations in maize tissue can inform breeding strategies and help develop more resilient short-stature varieties that maintain desired height characteristics across diverse growing conditions.

Understanding the natural range and stability of GA levels in conventional hybrid maize across diverse environments becomes essential for contextualizing the GA changes in new hybrids and predicting their performance across different agricultural regions. Such comprehensive baseline data can inform both the development and deployment of GA-modified hybrids while providing insights into the inherent plasticity of GA regulation in field conditions. The purpose of this study is to quantify major bioactive endogenous GA_1_ and GA_4_ in maize tissues by LC-MS/MS across conventional maize hybrids, tissue types, and environments to provide a baseline understanding of the contribution of genetics and environment to the variation of GA content to suggest potential targets for breeding programs aimed at improving agronomic traits such as yield.

## 2. Materials and methods

### 2.1. Maize field trials

Maize samples from ten commercially available conventional maize hybrids were collected from field trials during the 2023 growing season. Six field trial locations were spread across states representing major maize-growing regions of the United States (Iowa, Illinois, Indiana, Missouri, Nebraska, and Ohio) (Table 1)

**Table 1.** Field trial locations and maize hybrid names.

| Site Code | County, State | Hybrid Name |
| --- | --- | --- |
| Site 1 | Jefferson County, Iowa | Dekalb DKC58-32 |
|  |  | Lewis 12CV779 |
|  |  | Channel 210-79 |
| Site 2 | Shelby County, Illinois | Stone 6290 |
|  |  | Hubner H2390 |
|  |  | Dekalb DKC65-92 |
| Site 3 | Boone County, Indiana | Dekalb DKC63-58 |
|  |  | Hubner H2390 |
| Site 4 | Adair County, Missouri | Dekalb DKC58-32 |
|  |  | Lewis 12CV779 |
|  |  | Stone 6070 |
| Site 5 | York County, Nebraska | Stone 6290 |
|  |  | Hubner H2390 |
|  |  | Dekalb DKC64-32 |
| Site 6 | Miami County, Ohio | Dekalb DKC58-32 |
|  |  | Stewart 12CV209 |
|  |  | Channel 210-79 |

Each site was planted in a randomized complete block design with four replicated blocks. Normal agronomic practices were followed for each site. Maize samples were collected throughout the growing season and included the following tissues: leaf, root, stalk, pollen, forage, and grain. Sampling details and growth stage timing can be found in Table S1. All samples were kept on dry ice immediately after collection until storage in a -80°C freezer prior to analysis, with the exception of grain samples which were kept at ambient temperature.

### 2.2. Reagents

Gibberellin standards were used as reference standards for the analytical procedures or calibration of equipment. Stable-isotope-labelled gibberellin standards were used as internal standards (synthesized by OlChemlm). GA_1_ was synthesized by Toronto Research Chemicals (purity: 97%) and GA_4_ was synthesized by Thermo Scientific (purity: 90.30%). Formic acid (purity: 99%), ethyl acetate (HPLC grade) and methanol (purity: ≥99.8%) were purchased from Fisher Scientific and acetic acid (analytical grade) was obtained from Millipore, AR.

### 2.3. Sample preparation

After cryomilling maize tissues, 50 ± 5 mg of the sample was weighed into a pre-chilled 1.5 mL glass tube placed on dry ice. Next, on wet ice, 10 µL of internal standard working solutions (100 ng/mL of GA_1_-D4 and 25 ng/mL of GA_4_-D2) and 1.00 mL of pre-chilled extraction buffer (80% methanol with 0.5% formic acid) were added to each sample. The samples, arranged in a 96-well formatted rack, were manually agitated 40 times to thoroughly suspend the tissue in the extraction buffer, followed by an overnight incubation (approximately 15 hours) at 4°C with gentle mixing at 50 rpm on a Fisherbrand Nutating Mixter (Fisher Scientific, 88-861-043).After incubation, the samples were centrifuged at 2000 rpm for 10 minutes at 4°C. The supernatant was transferred to a 96-well polypropylene sample collection plate. The pellets were re-extracted by adding another 1 mL of extraction buffer following the same procedure as above. The combined supernatants evaporated under nitrogen flow at 35°C using a microplate sample evaporation system (Biotage SPE Dry 96, SD-9600-DHS-NA), until about 400 mL of aqueous phase was left. Subsequently, the extracts were loaded into a Biotage 96-well supported liquid extraction (SLE) plate (Biotage, 8200400P01), which was mounted on a glass-coated 96-well collection plate. The analyte GA was eluted with 2 × 0.9 mL of ethyl acetate. The eluate was then evaporated to dryness under nitrogen by using the SPE Dry 96. The dried samples were reconstituted in 100 µL of reconstitution buffer (20% Methanol with 0.1% formic acid), then vortexed at 1200 rpm at room temperature for 3 minutes. Following this, the samples were centrifuged at 2000 rpm for 3 minutes at 4°C, and 90 µL of the supernatant was transferred into a 96-well plate with glass inserts, kept on wet ice prior to LC-MS/MS analysis.

### 2.4. LC-MS/MS analysis of GA in maize tissues

Analytical instruments include mass spectrometer (Sciex triple Quad 7500), HPLC system (ExionLC 20A system) and data acquisition system (PC-based workstation with AB Sxiex OS software). Reconstituted samples were separated on a 1.7 µm Kinetex 100 Å C18 LC column (100 mm × 2.1 mm) (Phenomenex, 00D4475AN) at a flow rate of 0.40 mL/min and a column oven of 40°C. Mobile phase A is 0.5% acetic acid in water and mobile phase B is 0.5% acetic acid in methanol. The LC gradient is as below: mobile phase B (%) changes from 26 to 55 from 5.30 to 5.40 min, 55 to 58 from 5.40 to 10.20 min, 58 to 80 from 10.20 to 10.30 min, 80 to 81 from 10.30 to 12.70, 81 to 98 from 12.70 to 12.80, B (%) stays at 98 to 14.00 min, then drops from 98 to 26 from 14.00 to 14.10; finally, the column was equilibrated from 14.10 to 16 min to its initial conditions of 26% of B. Mass spectrometric conditions are as follows: polarity mode, negative ionization; scan type, MRM; spray voltage, -1800 V; ion source gas 1, 40 psi; ion source gas 2, 45 psi; source temperature, 500°C; curtain gas, 40; CAD gas, 10; entrance potential (EP), -10 V. The multiple reaction monitoring (MRM) transitions for each analyte and its stable isotope-labeled internal standard (IS) can be found in Table 2 with respective optimized collision energy (CE) and collision cell exit potential (CXP).

**Table 2.** MRM transition parameters.

| Compound ID | Q1 mass (Da) | Q3 mass (Da) | Dwell (ms) | CE(V) | CXP(V) |
| --- | --- | --- | --- | --- | --- |
| GA<sub>1</sub> | 347.19 | 273.101 | 100 | -32 | -8 |
| GA<sub>4</sub> | 331.19 | 213.131 | 100 | -40 | -7 |
| GA<sub>1</sub>-D4 (IS) | 351.19 | 277.069 | 100 | -36 | -9 |
| GA<sub>4</sub>-D2 (IS) | 333.14 | 259.064 | 100 | -33 | -9 |

### 2.5. MS data analysis

Raw data files were analyzed using Analytics in SCIEX OS software. A standard curve for each endogenous gibberellin was generated as the ratio of the gibberellin standard’s response (peak area) to the stable-isotope-labelled standard response (peak area), plotted against the concentrations of the gibberellin standard. A linear regression model was used for the quantification of the endogenous gibberellin levels with 1/x^2^ weighting. The final concentration of GA in the tissues was reported in pmol/g.

### 2.6. Variance component analysis to compare genetic and environmental variations

To compare how genetic and environmental factors contribute to expressions of GA_1_ and GA_4_ among five tissues, a variance component analysis [22] was performed with the following model (1) and a log transformation 𝑦 = log (𝑥), where 𝑥 is the observed response.

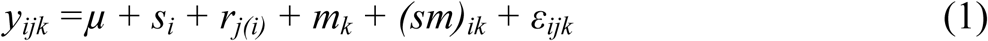

where *y_ijk_* is the observed response after transformation for the *k^th^* hybrid from the *j^th^* replicate at the *i^th^* site, *µ* is the overall mean, *s_i_* is the effect of the *i^th^* site, *r_j(i)_* is the effect of the *j^th^* replicate within the *i^th^* site, *m_k_*is the effect of the *k^th^* material, *(sm)_ik_* is the interaction effect of the *k*^th^ hybrid with the *i^th^* site, *ε_ijk_* is the residual. Before fitting the model, Levene’s tests were conducted to examine the assumption of the variance homogeneity, and non-significance was found for effects of sites, materials, and interactions on the variance. After fitting the model, the residual plots were examined for the normality assumption of the distribution. For the estimation of variance components, an all-random-effect model was assumed, and SAS procedure PROC MIXED was applied. Statistical significance of each component was determined by PROC GLIMMIX using a likelihood-based test at the 5% significance level. When the significance was confirmed, further investigation was made for the mean changes of individual hybrid across sites. In this case, a fixed-effect model for model (1) was assumed and the least square means were estimated. Notice that the fixed-effect model assumes the hybrid and site in the current experiment and the residual variation was the only source for the standard error [23]. All analyses were performed separately for each GA and each tissue, and results were presented in the following sections.

The statistical procedure SAS PROC MIXED was used to fit model (1) separately for GA_1_ and GA_4_. Residual plot showed no evidence of violation of normality and variance heterogeneity (as an additional visual confirmation to the above Levene’s test). Since the purpose of the analysis was to estimate the variation, all effects in model (1) were assumed to be random. The significance of each component was tested by the chi-square test with PROC GLIMMIX option COVTEST at the 5% significance level. The test is a likelihood-based method to account for the unbalanced design of the hybrid across sites (as explained further in discussion).

## 3. Results and discussion

### 3.1. Method development and validation of GA by LC-MS/MS

Plant tissue extracts contain complex mixtures and typically GA are present in trace amounts. Therefore, it is necessary to apply both a consistent sample preparation method to reliably extract GA from plant tissues and a sensitive technique to accurately quantify GA in plants. There has been continuous research on the quantitative analysis of GA in different crops[24]: the endogenous levels of eight GA were determined in beechnuts by GA chromatography-mass spectrometry[25]; 20 GA were analyzed as free acids by ultra performance liquid chromatography-tandem mass spectrometry[26] (UPLC-MS/MS) with plant extracts of *Brassica napus* and *Arabidopsis thaliana*; a phytohormone profiling method was established for rice[27]; and a high-efficiency method was developed for simultaneous quantitation of bioactive GA in *Litchi chinensis* using UHPLC-Triple Quadrupole--MS/MS[16]. However, the quantification of GA in different maize tissues has been less studied, with limited publications documenting their levels across different growth environments. For example, studies have primarily focused on the effects of exogenous GA applications on maize growth rather than on the natural variability of endogenous GA levels[28]. This gap highlights the need for comprehensive studies to establish reference levels of GA in maize at various geographical locations, which is essential for understanding their role in plant physiology and optimizing agricultural practices.

In this study, A robust, reliable and high-throughput method was developed for the simultaneous quantitation of GA in six maize tissue types (grain, pollen, leaf, root, forage, and stalk). In brief, the GA was extracted from approximately 50 mg of sample, purified by SLE, and separated and quantified by LC-MS/MS using the isotope-labeled internal standard (IS) as shown in Fig 1. The method was validated by determining the linearity (R^2^ ≥ 0.98) over the concentration range of 20-30,000 pg/mL in six tissue types. 20 pg/mL is the LLOQ (lower limit of quantification). Precision of the method was obtained with intra-and inter-day relative standards deviations (≤17.13%) and accuracy for compounds typically ranging between 79-124%. The recoveries of our method ranged from 77.6% to 119.8% in six maize tissues (Table 3). The bench-top stability of the tissue extraction supernatant was evaluated at room temperature for each type of tissue, demonstrating stability for a duration of up to two hours for all tissues except pollen, which exhibited stability of only one hour. Additionally, the stability of the processed samples in the autosampler was assessed at a temperature range of 4-8°C, with all tissue samples maintaining stability for up to six days.

**Fig 1.**
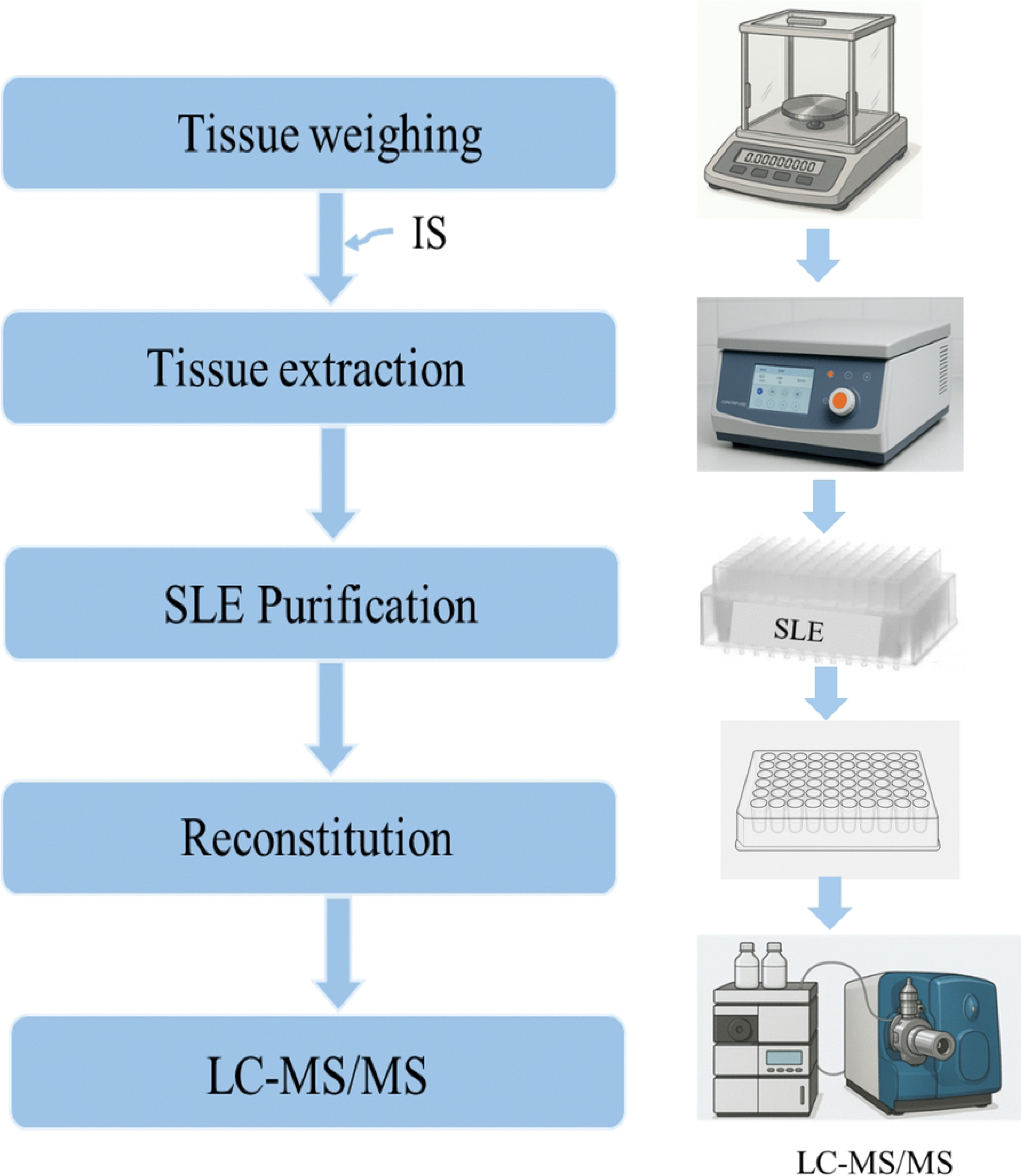
Workflow for Sample Preparation and Analysis of Gibberellins in Maize Tissues. Left Panel: The workflow consists of 5 key steps: (1) sample preparation via cryomilling and weighting, (2) extraction of gibberellins with internal standards (IS) and extraction buffer, (3) SLE analyte purification through centrifugation and supernatant collection, (4) Reconstitution of the analyte, and (5) LC-MS/MS analysis for quantitation. Right Panel: Equipment and components used in the workflow include a balance for weighing samples, a centrifuge for supernatant separation, a SLE plate for sample purification, a 96-well plate for sample collection and reconstitution, and an LC-MS/MS system for gibberellin quantification.

**Table 3.** Accuracy, precision, extraction efficiency and recovery for six maize tissues.

| Tissue type | Analytes | Accuracy (%) |  | Precision (CV%) |  | Extraction Efficiency (%) | Recovery (%) |
| --- | --- | --- | --- | --- | --- | --- | --- |
|  |  | Intra-assay | Inter-assay | Intra-assay | Inter-assay |  |  |
| Stalk | GA<sub>1</sub> | 83.92-114.86 | 95.78-100.71 | 0.56-11.5 | 1.80-14.52 | 94.56 | 96.56 |
|  | GA<sub>4</sub> | 87.69-113.10 | 90.62-105.81 | 0.42-11.69 | 2.51-9.33 | 91.52 | 115.72 |
| Grain | GA<sub>1</sub> | 79.73-115.95 | 89.38-106.18 | 0.49-12.39 | 1.65-17.13 | 95.91 | 92.01 |
|  | GA<sub>4</sub> | 85.83-110.13 | 86.33-108.89 | 0.99-12.52 | 1.48-11.78 | 93.02 | 77.59 |
| Leaf | GA<sub>1</sub> | 89.03-115.97 | 98.55-107.07 | 0.32-11.38 | 2.21-15.88 | 93.67 | 82.59 |
|  | GA<sub>4</sub> | 82.38-108.36 | 90.10-105.68 | 0.53-12.52 | 2.16-10.61 | 91.39 | 82.33 |
| Root | GA<sub>1</sub> | 83.92-114.86 | 95.78-104.13 | 0.24-11.50 | 1.80-14.52 | 96.51 | 98.52 |
|  | GA<sub>4</sub> | 85.23-108.36 | 90.62-103.43 | 0.42-11.69 | 2.51-9.43 | 91.08 | 107.03 |
| **Forage** | GA<sub>1</sub> | 83.92-123.06 | 92.36-110.33 | 0.32-9.13 | 2.81-13.98 | 94.44 | 93.40 |
|  | GA<sub>4</sub> | 87.69-108.95 | 90.62-103.43 | 0.42-11.69 | 2.51-9.33 | 91.71 | 101.89 |
| **Pollen** | GA<sub>1</sub> | 79.79-115.95 | 95.18-105.44 | 0.32-11.38 | 2.21-14.28 | 94.78 | 86.95 |
|  | GA<sub>4</sub> | 81.63-110.13 | 88.31-108.89 | 0.99-12.52 | 1.48-11.78 | 91.70 | 76.89 |

### 3.2. Quantification of GA in different maize tissue samples

The GA in maize samples were extracted, purified, separated and quantified by LC-MS/MS. Due to the short pollen tissue stability time, which was insufficient for handling the large number of samples, pollen data was not reported in this study. In total 68 individual samples of each of the five other tissue types were analyzed and reported. Data with non-detection (ND) values were assigned values by replacing ND with half of the observed minimums for that tissue in order to apply the statistical analysis. The details regarding values given to ND samples are included in Table S2. GA concentrations in maize tissues were calculated based on the standard curve and the statistics were summarized in Table 4.

**Table 4.** Summary of GA_1_ and GA_4_ concentration in different maize tissues.

| Analyte | Tissue | Mean (pmol/g) | Standard Error | Min (pmol/g) | Max (pmol/g) |
| --- | --- | --- | --- | --- | --- |
| GA<sub>1</sub> | Forage | 7.9 | 2.61 | 0.4 | 131.1 |
| GA<sub>1</sub> | Grain | 1.4 | 0.43 | 0.1 | 26.0 |
| GA<sub>1</sub> | Leaf | 3.2 | 0.57 | 0.4 | 27.0 |
| GA<sub>1</sub> | Root | 0.5 | 0.04 | 0.1 | 1.3 |
| GA<sub>1</sub> | Stalk | 2.1 | 0.08 | 0.9 | 3.4 |
| GA<sub>4</sub> | Forage | 1.9 | 0.56 | 0.1 | 29.2 |
| GA<sub>4</sub> | Grain | 0.9 | 0.06 | 0.3 | 2.0 |
| GA<sub>4</sub> | Leaf | 0.3 | 0.03 | 0.1 | 1.4 |

| Analyte | Tissue | Mean<br>(pmol/g) | Standard<br>Error | Min<br>(pmol/g) | Max<br>(pmol/g) |
| --- | --- | --- | --- | --- | --- |
| GA<sub>4</sub> | Root | 0.6 | 0.03 | 0.1 | 1.8 |
| GA<sub>4</sub> | Stalk | 0.4 | 0.02 | 0.2 | 0.9 |

This study provides quantification of endogenous GA_1_ and GA_4_ across five maize tissues collected from diverse field environments across the United States. Among these tissues, forage exhibited the highest GA concentrations and the broadest dynamic range, with GA_1_ ranging from 0.4 to 131.1 pmol/g and GA_4_ from 0.1 to 29.2 pmol/g. While there is limited study for forage, the shoot value has been studied. GA_1_ levels in shoot (used as a proxy for forage) ranged from 0.04 to 274.4 pmol/g [29], [30]. GA_4_ levels in the shoot tips range from 0.06 to 1.05 pmol/g [17] [31], which are notably lower than the maximum forage GA_4_ values observed in this study. This suggests that field-grown forage may accumulate higher GA_4_ levels, potentially due to environmental stimulation or hybrid-specific biosynthetic activity.

Stalk showed a narrower and more stable range (GA_1_: 0.9–3.4 pmol/g; GA_4_: 0.2–0.9 pmol/g), aligning closely with literature values (GA_1_ ∼1.8 pmol/g; GA_4_: 0.33–0.98 pmol/g) [15]. This consistency supports the use of stalk as a stable reference tissue for GA quantification, likely due to its structural maturity and reduced metabolic flexibility.

Leaf in this study showed GA_1_ levels ranging from 0.4 to 27.0 pmol/g and GA_4_ from 0.1 to 1.4 pmol/g. The GA_1_ concentrations observed here are higher than literature-reported ranges of 0–1.8 pmol/g, possibly due to the limited number of samples reported previously, while GA_4_ levels fall within the broader literature range of 0–3.5 pmol/g [15]; [32]; [31]. The elevated GA_1_ levels may reflect enhanced biosynthesis in field-grown hybrids or differences in developmental stage at sampling. In contrast, the GA_4_ levels, although lower than the upper literature limit, are consistent with expected physiological ranges and suggest that GA_4_ biosynthesis in leaf tissue may be more tightly regulated or less variable than GA_1_.

Grain exhibited GA_1_ levels from 0.1 to 26.0 pmol/g and GA_4_ from 0.3 to 2.0 pmol/g, exceeding literature ranges (GA_1_: 0.29–2.0 pmol/g; GA_4_: 0–1.2 pmol/g) [19] [33]. This suggests that grain GA levels may be more variable than previously reported. The tight range in previous literature may be due to the limited number of the samples analyzed or due to hybrid-specific traits or environmental modulation during seed development [33].

Root in this study showed GA_1_ concentrations ranging from 0.1 to 1.3 pmol/g and GA_4_ from 0.1 to 1.8 pmol/g. While literature data for GA levels in maize roots are limited or not reported, these findings provide a valuable baseline for future studies. The relatively narrow range and lower mean values suggest that GA biosynthesis in root tissues may be tightly regulated or less responsive to environmental variation compared to aboveground tissues. These results also highlight the importance of including below-ground tissues in hormonal profiling to better understand whole-plant hormone dynamics.

Overall, these comparisons highlight the broader dynamic range captured in this study, particularly for forage, leaf, and grain tissues. The elevated GA levels observed in several tissues may be linked to heterosis, as previous studies have shown that F1 hybrids often exhibit enhanced GA biosynthesis [29]. The comparison of literature data with our findings reinforces the robustness of the LC-MS/MS method and provides a valuable reference for future studies on GA-mediated growth regulation in maize. The observed tissue-specific and environmentally responsive patterns of GA accumulation offer critical insights for crop development initiatives focused on optimizing plant stature, biomass, and stress resilience.

### 3.3. Variance components in five tissues

Estimates of variance components for five tissues are listed in Table 5. The model variance was calculated as the sum of all components except the residual. Results in Table 5, in combination with means in Table 4 demonstrate large differences in distributions of GA_1_ and GA_4_ among the five tissues. For example, forage has the largest means as well as the largest residual variances for both GA_1_ and GA_4_, while stalk has intermediate means but the lowest residual variances. However, stalk shows higher model variation in percentage than forage (as shown in Fig 2). Out of ten (GA by tissues) combinations, only GA_4_ in grain shows significant hybrid contribution (Varg), as compared with five significant site contributions (Vars). To count all significant contributions, six out of ten GA and tissue combinations show at least one significant model component which is either site, hybrid, or hybrid by site interaction (Vargxs). The credibility of the data was confirmed, as only root has no significant components for both GA_1_ and GA_4_ among the five tissues. Also noticeably, the hybrid by site interaction was significant for both GA_1_ and GA_4_ in stalk.

**Fig 2.**
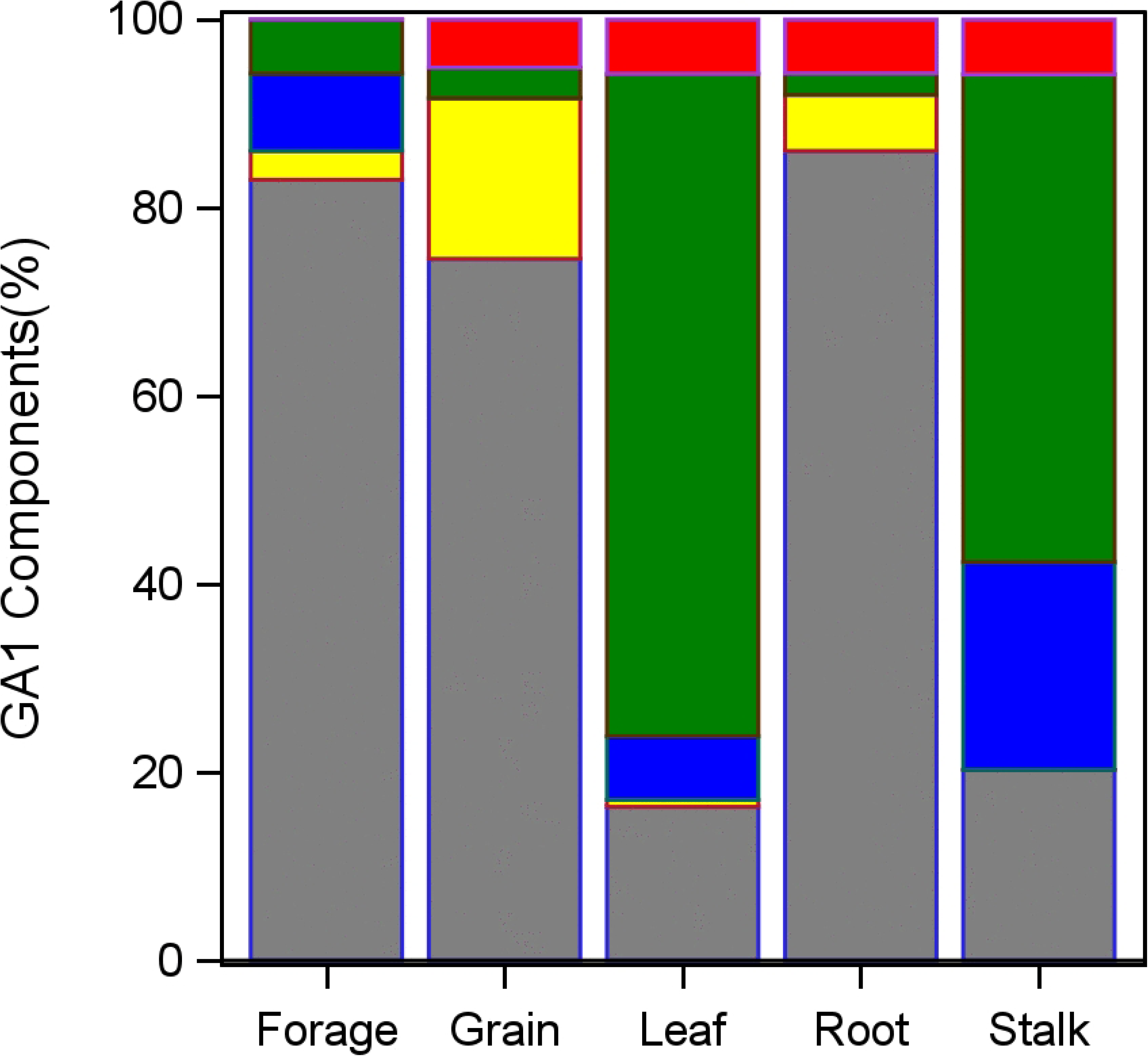
Variance Component Analysis of Genetic and Environmental Contributions to GA_1_ and GA_4_ Regulation. The figure presents the variance component analysis of gibberellin A1 (GA_1_) and gibberellin A4 (GA_4_) levels across different tissues in maize, broken down into contributing sources of variation. Bar charts show the percentage contributions of variance components: Varg (genetic variance, red), Vars (site variance, green), Vargxs (genetic-by-site interaction variance, blue), Varr (replicate variance, yellow), and Vare (residual variance, gray).(A) Variance components for GA_1_(B) Variance components for GA_4_

**Table 5.** Estimated variance components in log scale and the statistical significance from each source.

| Analyte | Tissue | Variance<br>(Model-log) | Variance Components (log)<sup>1</sup> |  |  |  |  |
| --- | --- | --- | --- | --- | --- | --- | --- |
|  |  |  | Varg | Vars | Vargxs | Varr | Vare |
| GA<sub>1</sub> | Forage | 0.39 | 0.00 | 0.13 | 0.19 | 0.07 | 1.88 |
| GA<sub>1</sub> | Grain | 0.41 | 0.08 | 0.05 | 0.00 | 0.28 | 1.21 |
| GA<sub>1</sub> | Leaf | 1.29 | 0.09 | 1.09* | 0.10 | 0.01 | 0.25 |
| GA<sub>1</sub> | Root | 0.11 | 0.04 | 0.02 | 0.00 | 0.05 | 0.67 |
| GA<sub>1</sub> | Stalk | 0.10 | 0.01 | 0.07* | 0.03* | 0.00 | 0.03 |
| GA<sub>4</sub> | Forage | 0.73 | 0.00 | 0.58* | 0.00 | 0.14 | 1.09 |
| GA<sub>4</sub> | Grain | 0.19 | 0.06* | 0.09* | 0.01 | 0.02 | 0.07 |
| GA<sub>4</sub> | Leaf | 0.65 | 0.05 | 0.54* | 0.07 | 0.00 | 0.47 |
| GA<sub>4</sub> | Root | 0.05 | 0.00 | 0.01 | 0.03 | 0.01 | 0.20 |
| GA<sub>4</sub> | Stalk | 0.05 | 0.00 | 0.00 | 0.05* | 0.00 | 0.05 |

<sup>1</sup> hybrid (Varg), site (Vars), hybrid by site interaction (Vargxs), replicate (Varr), and residual (Vare). Variance (Model) is the sum of all components except Vare.
\* indicates statistical significance at the 5% level.

The relative contribution of variance components in terms of the percentage of each source to the total variance is presented in Fig 2. As shown in Table 5 and Fig 2, the residual variation (Vare) is greater than the model variation in five out of ten combinations. Regardless of the large residual variation, the variance components in Table 5 and Fig 2 were estimated by an all-random-effect model, and any statistical significance represents the chance if the experiment and the analysis were repeated in the same design with different sets of sites and hybrids.

In this investigation, the contribution of different sources of variance components to the expression was analyzed by an all-random-effect model. The model (1) includes effects of site, hybrid, hybrid by site interaction, replicate, and residual. The hybrid effect represents the genetic effect, site effect may include various environmental factors, and the residual includes the plant sample variation in the field and the assay variation in the lab. Results in Table 5 and Fig 2 indicated that while both genetic and environment are important factors, the site is a major contributor as compared with the genetic background to the expression. In addition, in five out of ten (GA by tissue) combinations, residual variations are greater than the model variation especially for forage and root.

### 3.4. Hybrid by site interaction of GA_1_ and GA_4_ in leaf and stalk

The significant hybrid by site interactions for both GA_1_ and GA_4_ levels in stalk in Table 5 is especially noticeable. To visualize the interaction, hybrid means at each of six sites were estimated by the least square means under a fixed-effect model as explained in the previous section, and then back-transformed into the original unit in pg/mL along with their standard errors. These hybrid means were plotted across six sites in Fig 3. Three hybrids were planted at each site, with the exception of site 3 with two hybrids. Out of ten hybrids, five were planted only at a single site, the remaining five were each from two or three sites and connected by lines across corresponding sites. Six sites were ordered by the magnitude of the site effects of GA_1_ from low to high. In contrast, the same plot was created for leaf which showed significant site variations for both GA_1_ and GA_4_ in Table 5 but non-significant hybrid by site interactions. The hybrid means of leaf show a clear decline over sites for both GA_1_ and GA_4_ but much smaller difference among hybrid means averaged over sites, which is in line with the significant site component and non-significant hybrid component in Table 5. The decline trend over sites in leaf is in contrast with the increasing trend in stalk. For stalk, lines of four hybrids for GA_1_ and three for GA_4_ were approximately parallel over sites, that is, the expression was consistent over sites with no evidence of interaction. But the hybrid Dekalb DKC58-32 shows a different trend with the highest expressions at site 4 for both GA_1_ and GA_4_, and reduced expression at site 1 where other hybrids tend to increase the expression (except for GA_4_ of Channel 210-79). Consistent trends in means of GA_1_ and GA_4_ contribute to a positive correlation between the two, as discussed in the following section. By comparing the error bars with the hybrid mean changes over sites, the significant site variation for leaf and the significant interaction for stalk were visually confirmed.

**Fig 3.**
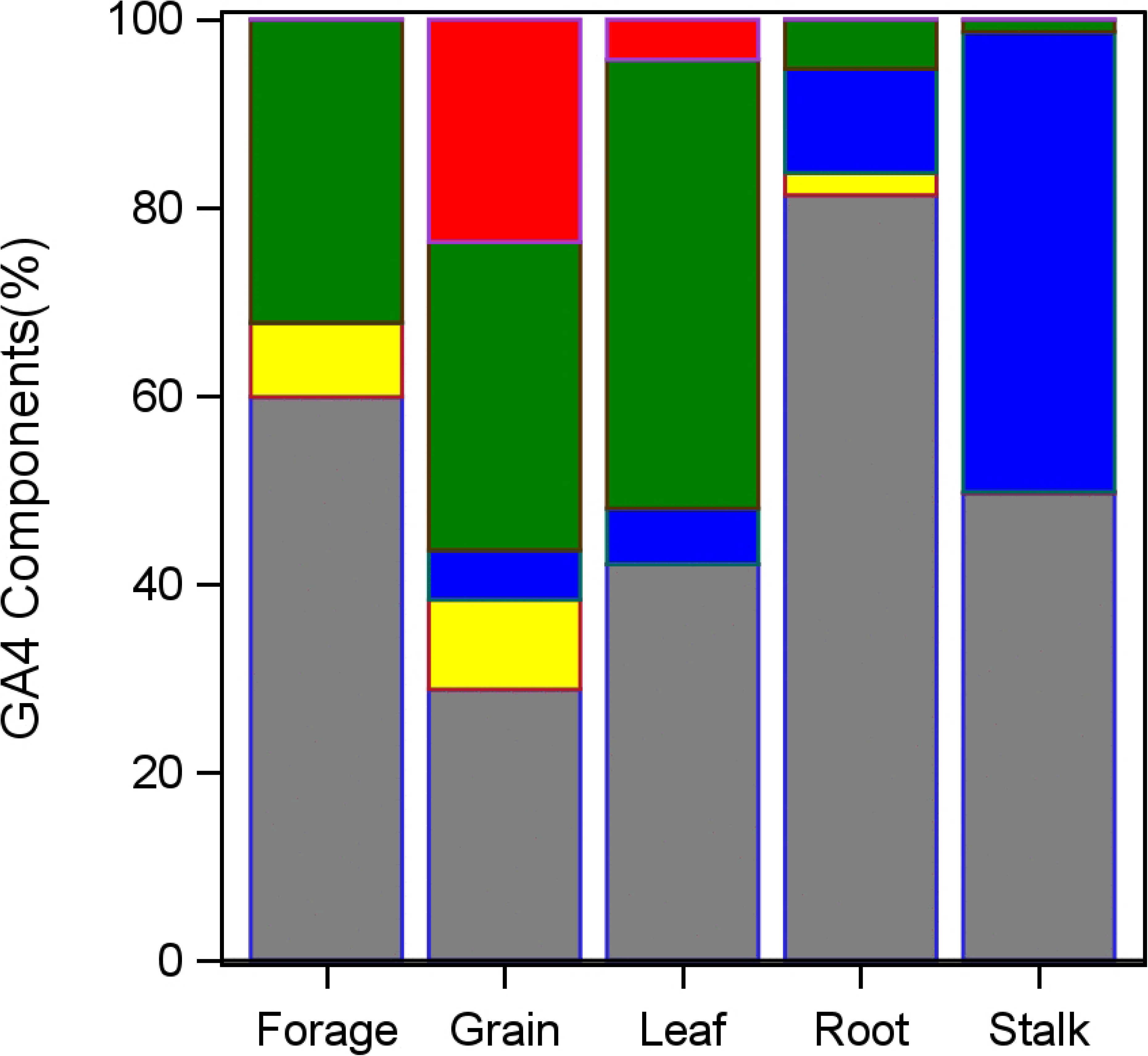
GA_1_ and GA_4_ Expression in Leaf and Stalk Across Hybrids and Sites. Expression levels of gibberellin A1 (GA_1_) and gibberellin A4 (GA_4_) (mean ± SE) in leaf and stalk were measured and plotted for 10 maize hybrids across six field sites. Data are presented in pmol/g, with hybrids represented by different markers and colors, and site-wise trends connected by lines for visualization **Panels:** Top Left: Leaf GA_1_ expression, Top Right: Leaf GA_4_ expression, Bottom Left: Stalk GA_1_ expression, Bottom Right: Stalk GA_4_ expression

The design of this experiment was limited by factors such as hybrid adaptation and site capacity, resulting in selected genetic backgrounds (i.e. hybrids) for each site. This unbalanced arrangement is expected to impact the assessment (e.g. the statistical significance) of the variance components. For example, the experiment included ten hybrids and six sites. With a balanced design with all hybrids at each site, the interaction degrees of freedom would have been 45. However, the mixed model analysis of the available data showed only three degrees of freedom for interaction. To ensure the repeatability of the conclusion from this analysis, the variance component analysis assumed an all-random-effect model and the estimated components and their significance from the analysis are expected to be unbiased by the current design. As one of the main conclusions shown in Fig 3, GA expressions of a large proportion of genetic backgrounds appear to be parallel across sites, however, some genetic backgrounds may show both genetic and environmental contributions are important regardless of the non-significance of main effects of the hybrid and the site in this experiment. For example, the hybrid Dekalb DKC58-32 showed the highest GA expression among all hybrids at site 4 but relatively lower expression at other sites. Despite the environmental cause of the highest GA expression of this hybrid-site combination cannot be confirmed, the dependence of GA expression on environment is clearly shown. Factors such as temperature, humidity, and rainfall are all expected to be relevant, and further experimentation is necessary to show their relative importance.

### 3.5. Correlation between GA_1_ and GA_4_ across tissues

The correlation between GA_1_ and GA_4_ was investigated using the biplots. The plot is based on the hybrid mean at each site as shown in Fig 4. In this plot, each mark represents the observed predicted hybrid mean over four replicates in log and then back transformed into pmol/g; five tissues are represented by different markers and colors. As shown in Fig 4, positive correlations were observed in four out of five tissues except root. The position and range of each tissue are represented by the differences in means (see Table 4) and the variation (see Table 5) among tissues. Since the hybrid mean at each site in Fig 4 consists of both the hybrid and site effects on the expressions, these positive correlations are due to similar trends in hybrid mean changes over sites between two GA as shown in Fig 3. The regression coefficients of GA_4_ on GA_1_ in log were estimated as 0.01, 0.42, 0.42, 0.40, 0.49 for root, forage, leaf, grain, stalk with P values of 0.94, 0.10, 0.03, <0.01, <0.01, respectively, which are significant at the 5% level for leaf, grain, and stalk, and non-significance for forage.

**Fig 4.**
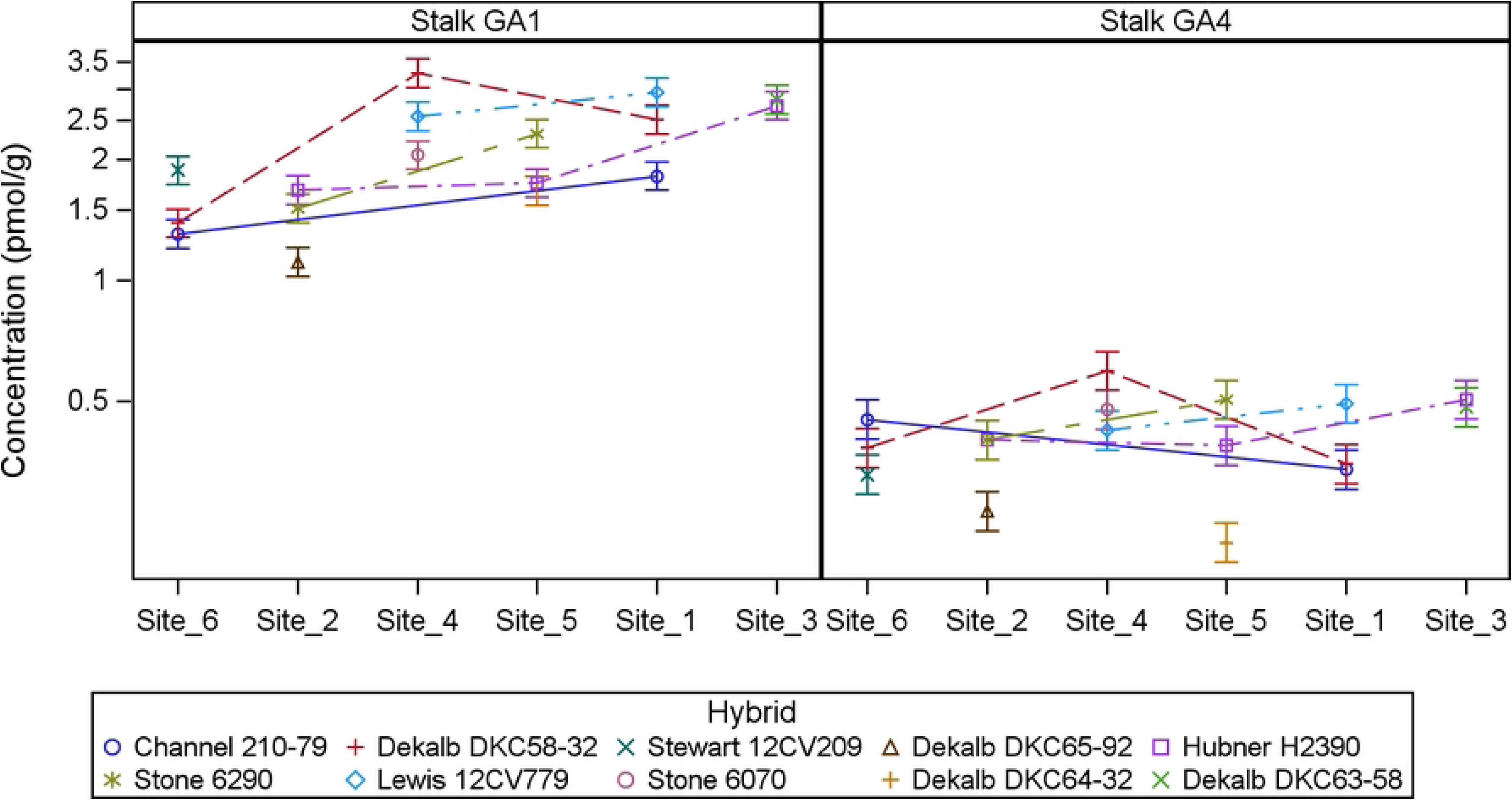
Biplot and Correlation Between GA_1_ and GA_4_ Means Across Five Maize Tissues. The figure illustrates the relationship between gibberellin A1 (GA_1_) and gibberellin A4 (GA_4_) concentrations across five maize tissues: root, forage, leaf, grain, and stalk. Each marker in the biplot represents the predicted hybrid mean for GA_1_ and GA_4_ at a specific site, derived from four replicates, expressed in pmol/g (log-transformed and back-transformed). Different tissues are distinguished by unique markers and colors.

In this study, a positive correlation was observed between GA_1_ and GA_4_ across four different tissues in maize hybrids, with the exception of roots. The correlation is consistent on the level of the hybrid mean changing over sites as well as the level of individual sample’s deviation from the mean. This correlation is particularly intriguing given that GA_1_ and GA_4_, despite being structurally related and bioactive GA, are synthesized through distinct hydroxylation pathways. Both GA are known to play critical roles in regulating plant growth and development at various stages. The correlation between GA_1_ and GA_4_ suggests potential shared regulatory mechanisms or environmental factors influencing their biosynthesis, warranting further investigation into the underlying genetic and biochemical pathways.

These findings highlight the complexity of hormonal regulation in plant growth, suggesting that a multifaceted approach is necessary to fully understand the interplay between different hormones and external conditions. Further research is needed to elucidate the mechanisms driving these interactions and their impact on plant development.

## 4. Conclusions

In this study, a validated high-throughput LC-MS/MS method has been applied to simultaneously quantify the bioactive gibberellins GA_1_ and GA_4_ across multiple tissues of commercial maize hybrids grown in North American field environments. Our work provides the first comprehensive baseline of natural GA_1_ and GA_4_ variation under real-world agricultural conditions. We found that GA concentrations varied significantly across tissues, with forage exhibiting the broadest dynamic range and highest levels, while stalk tissue showed more stable and narrow concentration ranges, suggesting its utility as a reference tissue for future studies.

A key finding of this research is the profound influence of the environment on gibberellin regulation. Variance component analysis revealed that the field site was the most significant factor driving the variability in GA_1_ and GA_4_ levels for most tissues. This highlights the remarkable plasticity of the gibberellin pathway in response to local environmental conditions and underscores the importance of considering environmental context in studies of hormonal regulation. Critically, we identified a significant genetic (hybrid) contribution to GA_4_ variance specifically in grain. This novel insight suggests that GA_4_ levels in grain may be under a degree of genetic control, presenting a potential new target for hybrid selection and breeding programs aimed at optimizing agronomic traits such as yield.

Furthermore, our analysis uncovered a consistent positive correlation between GA_1_ and GA_4_ concentrations in forage, grain, leaf, and stalk tissues. This finding is particularly noteworthy as these two bioactive GA are synthesized via distinct downstream pathways. The observed co-regulation suggests the presence of a coordinated transcriptional network or shared upstream regulatory mechanisms that modulate both pathways in response to developmental and environmental cues.

While this study provides a foundational dataset, it is based on a single growing season. Future multi-year studies would help confirm the stability of these trends and further dissect the impact of annual weather variations. Nevertheless, the data and analyses presented here offer critical insights into the complex interplay between genetics, environment, and hormonal regulation in maize. This research establishes an essential framework for future investigations into the specific environmental drivers of GA expression and provides a compelling rationale for exploring the genetic basis of GA_4_ accumulation in grain as a potential avenue for crop improvement.

## Acknowledgments

The authors thank Leah Riter, Vinod Jakkula, Edward Cargill, Mohamed Bedair, David Caldwell, Prema Karunanithi and Paul Tietz for providing valuable feedback on the manuscript.

## Supporting information

**S1 Table.** Summary of sample type and collection information.

| Sample Type | Sampling Details | Collection Growth Stage | Growth Stage Description |
| --- | --- | --- | --- |
| Leaf | Approximately 15 cm of most recently collared leaf | V2-V4 | A plant with 2 to 4 leaf collars |
| Root | All below-ground root tissue, thoroughly washed | V2-V4 | A plant with 2 to 4 leaf collars |
| Stalk | All above-ground stalk tissue with leaf blade, leaf sheath, ear, and tassel removed | V10-V12 | A plant with 10 to 12 leaf collars |
| Pollen | Fresh pollen with all anthers removed | Pollen shed | A plant that is actively shedding pollen |
| Forage | All above-ground plant tissue | R5, early dent | Most kernels are dented and contain approximately 55% moisture |
| Grain | Shelled grain dried to $\leq 15.5\%$ moisture prior to shipping and storage | R6 | Physiological maturity; black layer visible at base of kernels |

**S2 Table.**
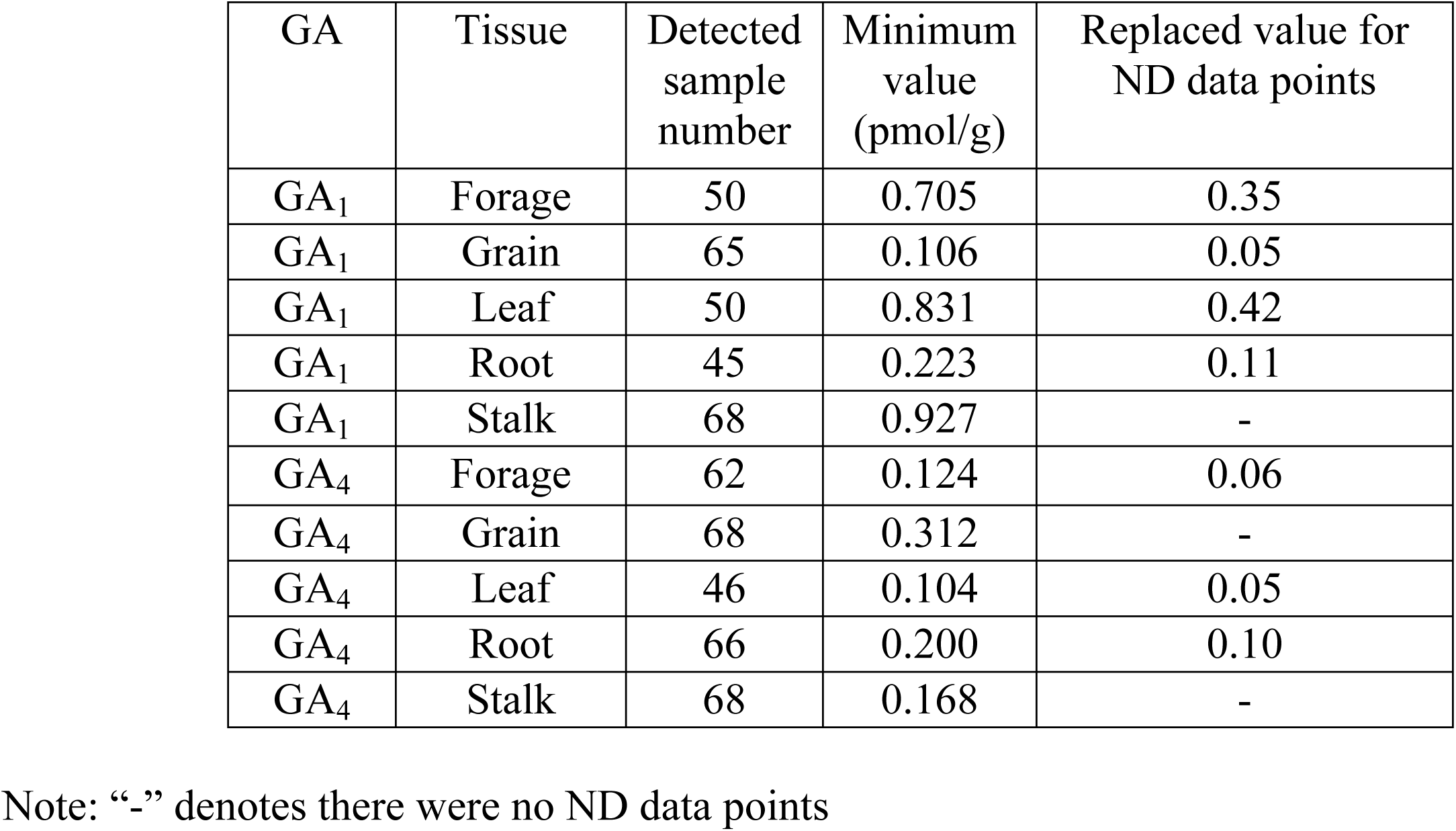
Value assigned to non-detection (ND) data points for statistical analysis.

## Author contributions

Credit: Yanfei Wang: Conceptualization, Project administration, Resources, Supervision, Writing – review & editing; Xia Guan: Formal analysis, Investigation, Methodology, Writing – original draft, Writing – review & editing; Nitin Supekar: Formal analysis, Investigation, Writing – original draft, Writing – review & editing; Changjian Jiang: Statistical analysis, Data curation, Formal analysis, Methodology, Writing – original draft, Writing – review & editing; Junhong Guo: Methodology, Formal analysis, Investigation, Writing – original draft, Writing – review & editing; Ed Allen: Project administration, Writing – review & editing; Charles Dietrich: Project administration, Writing – review & editing; Cody Postin: Formal analysis, Investigation, Methodology, Writing – original draft, Writing – review & editing;

---

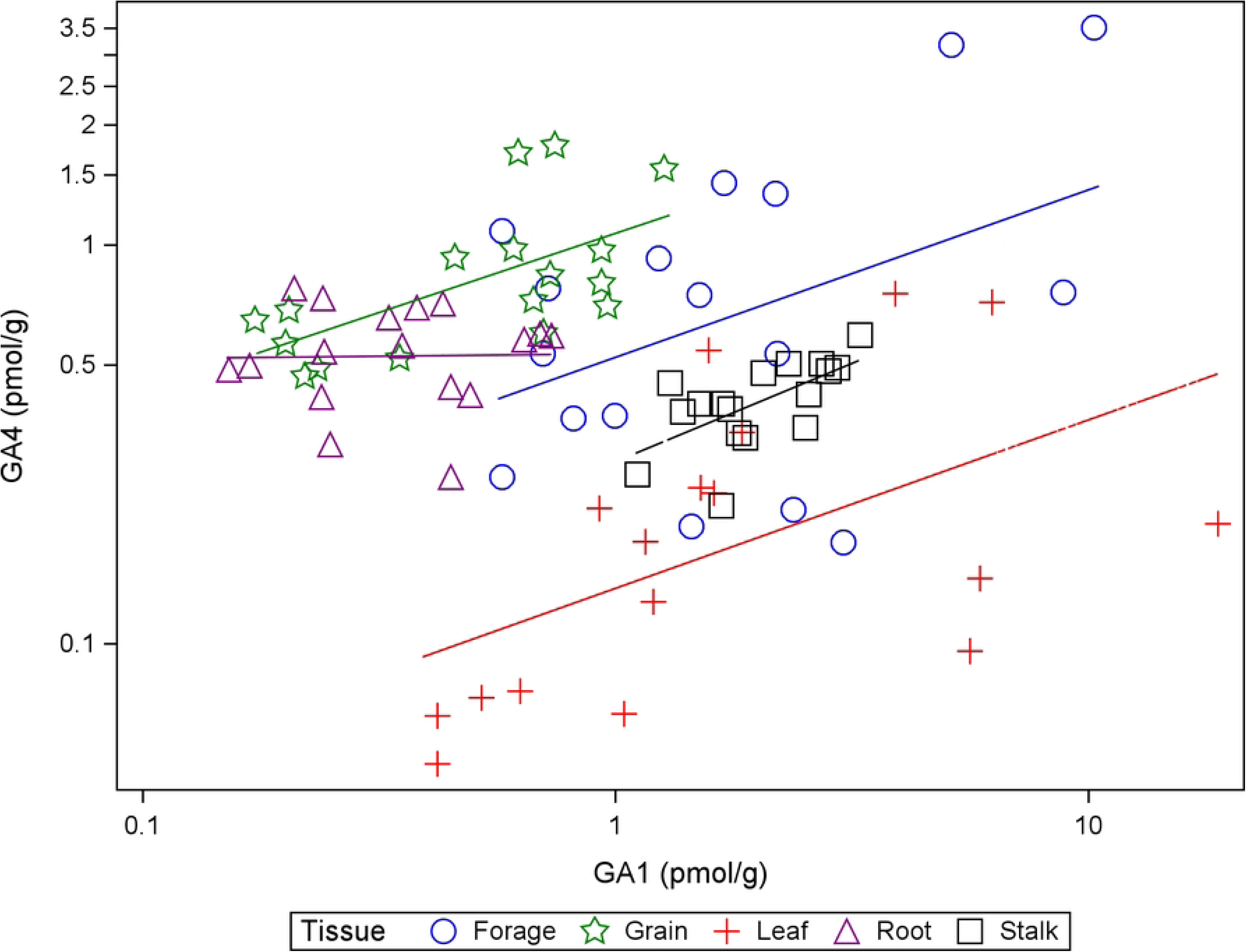

